# Systematic review of the validity of the hepatitis B virus detection tests in blood banks from 2000 to 2018

**DOI:** 10.1101/2020.04.17.046458

**Authors:** Claudia Patricia Orrego-Marín, Jaiberth Antonio Cardona-Arias, Sandra Medina-Moreno, Juan Carlos Zapata

## Abstract

**Background:** Hepatitis B Virus (HBV) is a public health problem that causes chronic hepatitis, eventually evolving to cirrhosis and hepatocellular carcinoma. Given the low frequency of screening in the general population, blood banks represent a key element for monitoring and controlling HBV transmission. The objective of this work was to evaluate the validity of the diagnosis of HBV in blood banks, based on studies published in the world scientific literature from 2000 to 2018.

**Methods:** We used a Meta-analysis of random effects with application of a search and selection protocol according to Cochrane and PRISMA guidelines. Reproducibility, completeness and quality assessment were affirmed with QUADAS, a tool for assessing diagnostics. The parameters of sensitivity, specificity, likelihood ratios, odds ratio and ROC curve were analyzed in MetaDisc with 95% confidence.

**Results:** From 4,061 studies screened, only 12 complied with the protocol, and the compliant studies included a population of 17,391 healthy people and 1,229 infected. The compliant studies evaluated mainly immunodiagnostics (ELISA) using HB surface antigen (HBsAg) and less frequently anti-HBc (HB core antigen) and PCR. The tests for HBsAg presented sensitivity of 94.1% (95% CI = 92.9% - 95.1%), specificity 98.2% (95% CI = 97.8% - 98.6%), diagnostic OR of 1721 (95% CI = 607.18 - 4418.8) and Area Under the Curve of 99.7%.

**Conclusion:** With a large sample size and with the high quality of the studies evaluated in this review, we confirm that HBsAg for HBV immunodiagnostics has excellent validity, which supports its use in clinical screening, blood banks and population surveillance programs.

## Introduction

Hepatitis B virus (HBV) remains one of the main public health problems worldwide and an important cause for the development of chronic hepatitis, cirrhosis and hepatocellular carcinoma [1-3]. It is estimated that about two billion people are serologically positive for HBV [4] and according to data from the World Health Organization (WHO) in 2017, approximately 325 million people suffer from chronic infection [1]; This situation is aggravated if it is considered that diseases such as cirrhosis and carcinoma cost more than 650,000 lives each year [2].

HBV transmission occurs through blood transfusion, direct contact with blood, sexual intercourse and/or through intravenous injections [2,5-7]. WHO estimated that in 2001, unsafe blood transfusions generated 8-16 million HBV infections [6]. Although the current treatment and vaccination options are helping to decrease this burden, especially the increased childhood vaccination coverage between 1980 and 2000 [1,4], the presence of mutations in the virus associated with selective pressure for vaccination and treatment, often makes it difficult to control this infection [2]. Furthermore, the poor availability of prevention programs, and the existence of individuals that do not respond adequately to the vaccine, means that almost 257 million people born before the modern vaccination regimes have an increased risk of infection. This situation is of even greater concern when considering the 2017 WHO global report that most infected people lack access to diagnosis, which means that there are millions of infected people at risk of developing chronic hepatitis, cancer and death, as well as being potential sources of the infection [1].

Given the problem of access to HBV diagnosis in some regions, blood banks represent a key element for the detection of infected people and for access to treatment. In order to assure high quality results, the application of technology with high diagnostic validity is required to avoid unnecessary deferral by false positives and the inherent risk in false negatives, especially those donors with chronic HBV infections and low viremia levels [4,6,8,9].

In this sense, detection of antigens and antibodies individually or in combination, is of great benefit for the timely diagnosis of HBV. The detection of the hepatitis B surface antigen (HBsAg) is the most common assay used by bloodbanks to check the infection; it appears earliest and can remain detectable for life [3, 9]. Also used is detection of antibodies against the hepatitis B core antigen (anti-HBc) that appears during the acute infection and signify a new infection [3,9].

Some previous studies have documented a high heterogeneity in the validity data of HBV screening tests. For example, Farooq *et al.*, in 2017 compared three routinely diagnostic tests for HBsAg (SD Bioline rapid assay, GB HBsAg ELISA and Abbott ARCHITECT^®^CLIA system) with a quantitative determination of hepatitis B surface antigen (HBsAg) (LIAISON^®^ XL Murex HBsAg Quant assay - chemiluminescence immunoassay), in which they found sensitivities of 17.2%, 43.7%, 90.9% and 100% respectively, and positive probability ratios between 38 and 100 [2]. In addition, Mutocheluh *et al.* in 2014 evaluated the performance of the most common immunochromatographic (ICT) kits (Wondfo, Rapid Care, Core TM, Accul-Tell and Abon) used for screening blood donors in some blood bank facilities in the northern part of Ghana. The reported sensitivities for the HBsAg rapid tests in this study were, Wondfo 59.1% Rapid Care and Accul-Tell 54.5%, and Core TM and Abon 50% [10]. On the other hand, Randrianirina *et al.*, in 2008 and Dogbe et *al.*, in 2015 reported overall sensitivities for rapid tests between 93.8% and 100%; with specificity values between 95.6% and 100% [4,6].

Those results presented above demonstrate a high diversity of diagnostic validity which underpins the need to meta-analyze the evidence available on this topic. In this sense, it should be considered that systematic reviews have advantages over the evidence of individual studies, by improving the external validity of the results, the quality of clinical and epidemiological recommendations, and the accuracy of the estimates. Therefore, it permits the generation of results from an exhaustive search of the literature containing studies of the same kind and with a common goal, allowing a combination of findings, while reducing biases and random errors of the reviewed literature. Particularly in diagnostic tests, systematic reviews allow for a more complete assessment by estimating parameters such as sensitivity, specificity, likelihood ratios (LR), diagnostic odds ratio (OR) and a receiver operating characteristic (ROC) curve [11-13]. Therefore, the objective of this research was to evaluate the validity of HBV immunodiagnosis, focusing on HBsAg, based on studies published in the scientific literature worldwide between 2000 and 2018.

## Materials and methods

### Study search and selection protocol

A systematic review of the literature with a meta-analysis of diagnostic tests was carried out following the recommendations of the Cochrane Collaboration and the four phases of the preferred reporting items for systematic reviews and meta-analyses (PRISMA) guide as described [14].

#### Identification

Specific search, circumscribed to language controlled in terms of thesaurus, particularly multilingual and structured vocabulary DeCS - Health Sciences Descriptors and MeSH - Medical Subject Headings, was performed for original articles published in PubMed, Scielo and Science Direct, which was supplemented by a sensitivity search on Google Scholar. Eight search terms were used for diagnostic evaluation parameters (false positives, false negatives, negative predictive value, positive predictive value, true negative, true positive, sensitivity and specificity), two synonyms for blood banks and blood banking, and two for blood donations. Each diagnostic parameter was combined with the four terms of the blood bank obtaining a total of 32 different search strategies (Supplementary Material 1). Some syntaxes used were: in PubMed *((Sensitivity[Title/Abstract]) AND Specificity[Title/Abstract]) AND Blood Donor[Title/Abstract] OR ((Sensitivity[Title/Abstract]) AND Specificity[Title/Abstract]) AND Blood Banks[Title/Abstract]*, in Science Direct: Title, abstract, keywords: Positive Predictive Value AND Blood Donation, in Scielo: ti:(ab:(Negative Predictive Value AND Blood Donation)))and in Google scholar (*allintitle: False Positives and Blood Donation)* (Supporting information 1. Search strategies applied).

#### Screening

Duplicate studies were eliminated, and two inclusion criteria were applied: original studies and blood donor research.

#### Eligibility

Studies were excluded that did not include HBV, or were not available in databases, or that failed to receive any response from the author. In addition, manuscripts with incomplete information (incomplete data on the variables to be analyzed in this review), or articles in which the tests would not be evaluated or did not have reports of sensitivity, specificity, positive or negative predictive values were all excluded from this study.

#### Inclusion

Those articles that fulfilled the previous criteria, were evaluated for methodological quality and extraction of variables. It includes title, lead author, publication year, place of study, population, type of diagnostic test analyzed, true positives (HBV-infected donors and positive screening result) and true negatives (donors without HBV with negative screening result), false positives (donors without HBV and positive screening result), and false negatives (donors with HBV with negative screening result).

### Reproducibility and quality assessment

To ensure reproducibility of the search and in the selection of studies, the search was carried out by two reviewers. It was determined *a priori* that the differences would be resolved by consensus. To ensure the reproducibility of the extraction of variables, two reviewers performed independently the search and the kappa index was applied for qualitative variables and intra-class correlation coefficient for quantitative variables. To assess the methodological quality of the studies included, the criteria of the Quality Assessment of Diagnostic Accuracy Studies (QUADAS) guide were applied [15].

### Analysis plan

The studies were described with absolute and relative frequencies. Sensitivity, specificity, positive and negative LR, diagnostic OR and ROC curve with their 95% confidence intervals were estimated. The analyses were performed with a random effects meta-analysis in *MetaDis (Meta analysis of studies of evaluations of Diagnostic and Screening tests)* with a significance of 0.05 and REM model. The combined measurements of the studies are based on the Q*(X*^2^) Der Simonian-Laird Test. 95% confidence intervals were corrected with an over-dispersion estimate. The heterogeneity analysis is based on Cochrane’s Q statistic and the uncertainty (sensitivity) statistic on the weight percentage of each individual study on the overall result.

## Results

4,601 studies were sifted with the search terms in title or abstract. From those only 12 complied with the search and selection protocol (Figure 1). Those studies were published between 2006 and 2017, most were conducted in African and Asian countries, in a total population of 18,620 subjects of which 17,391 were healthy and 1,229 were infected (Table 1).

**Table 1.**
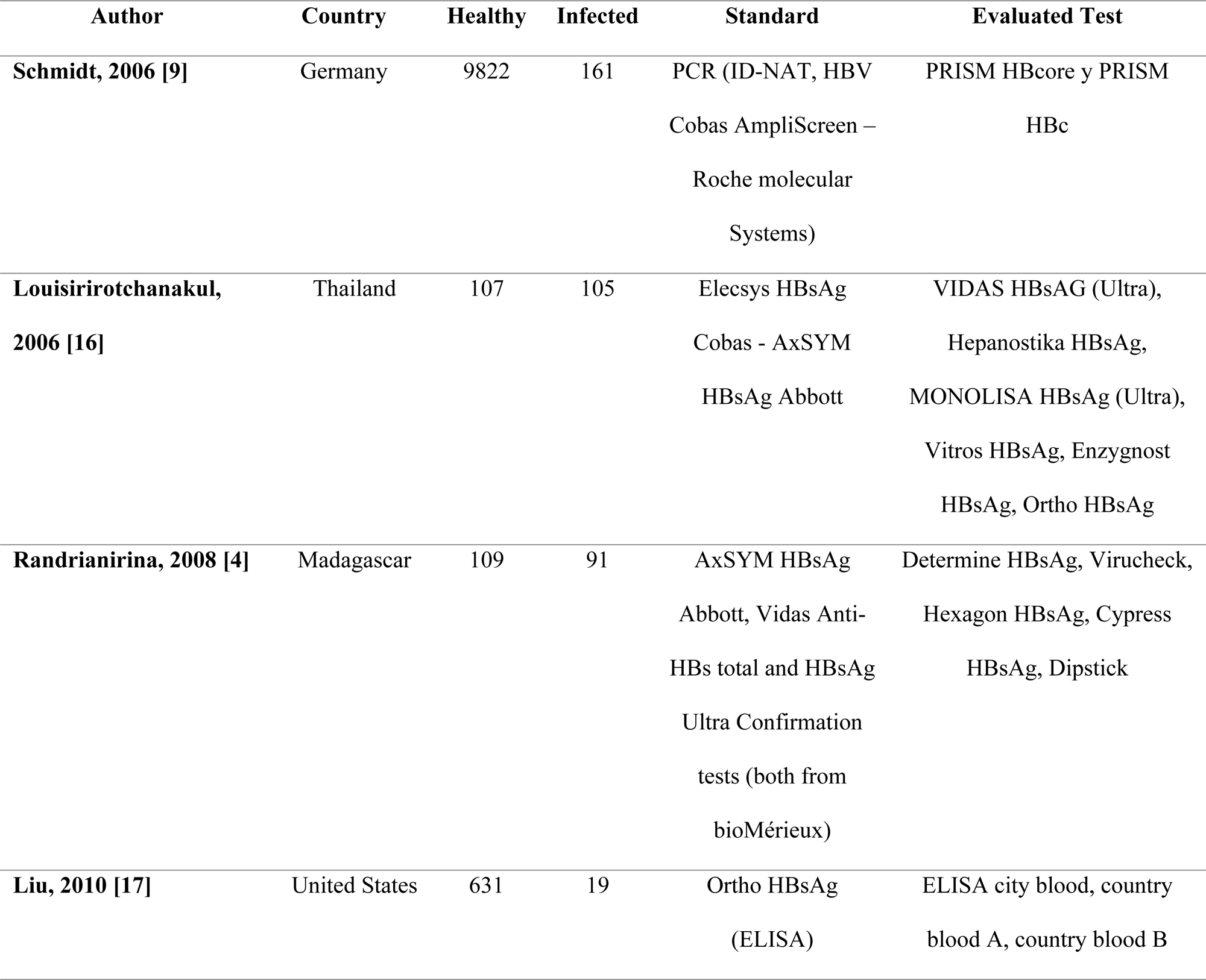
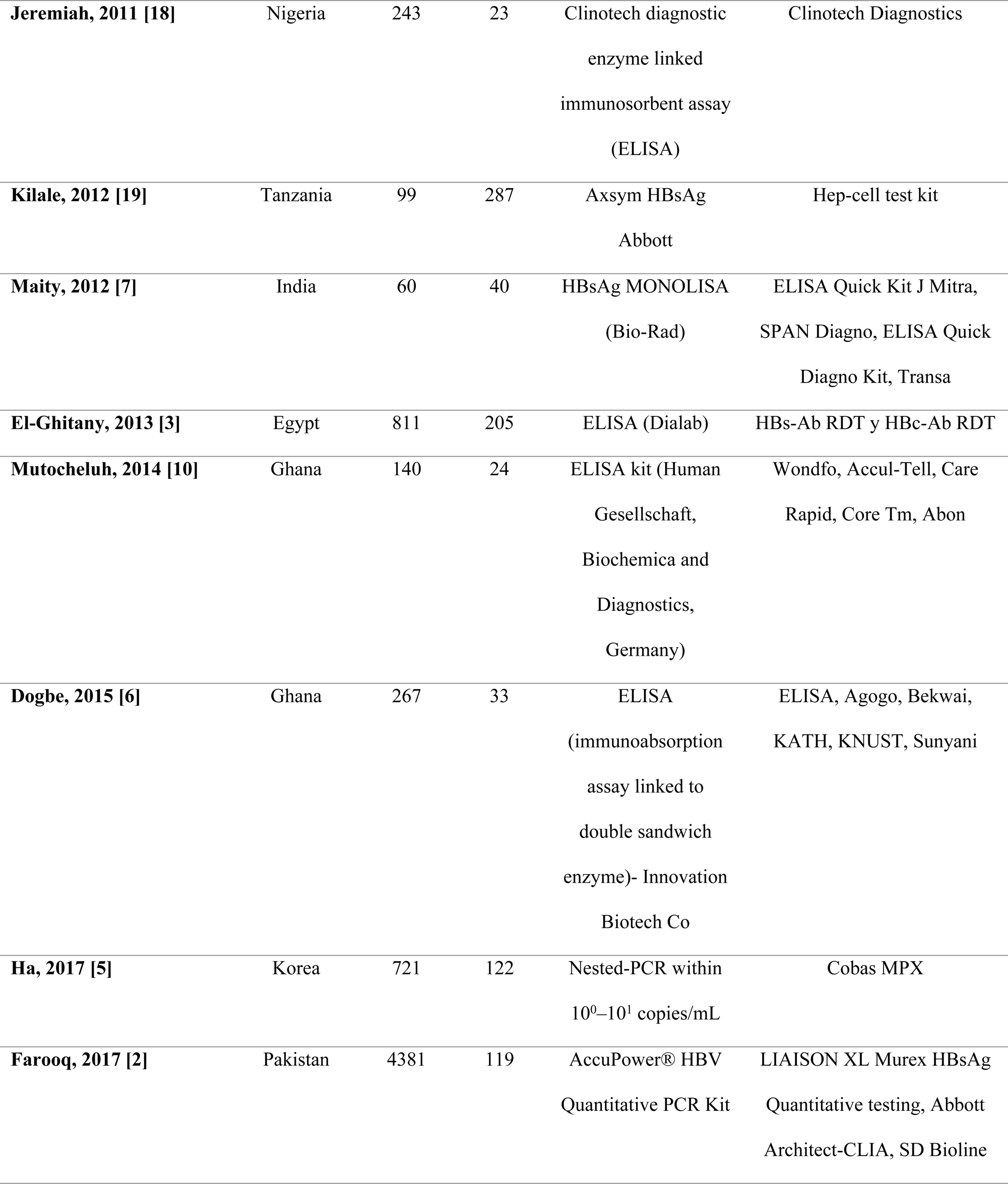
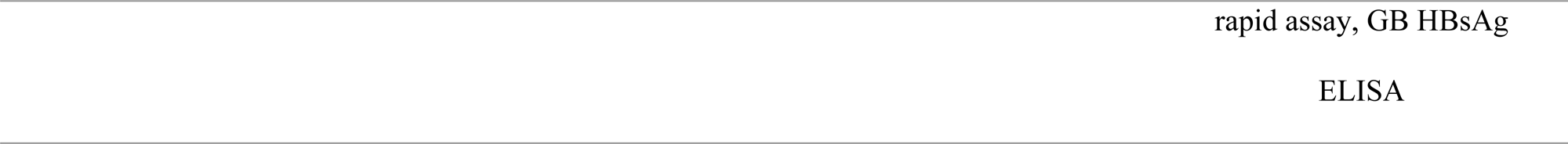
Description of the studies included in this review.

**Figure 1.**
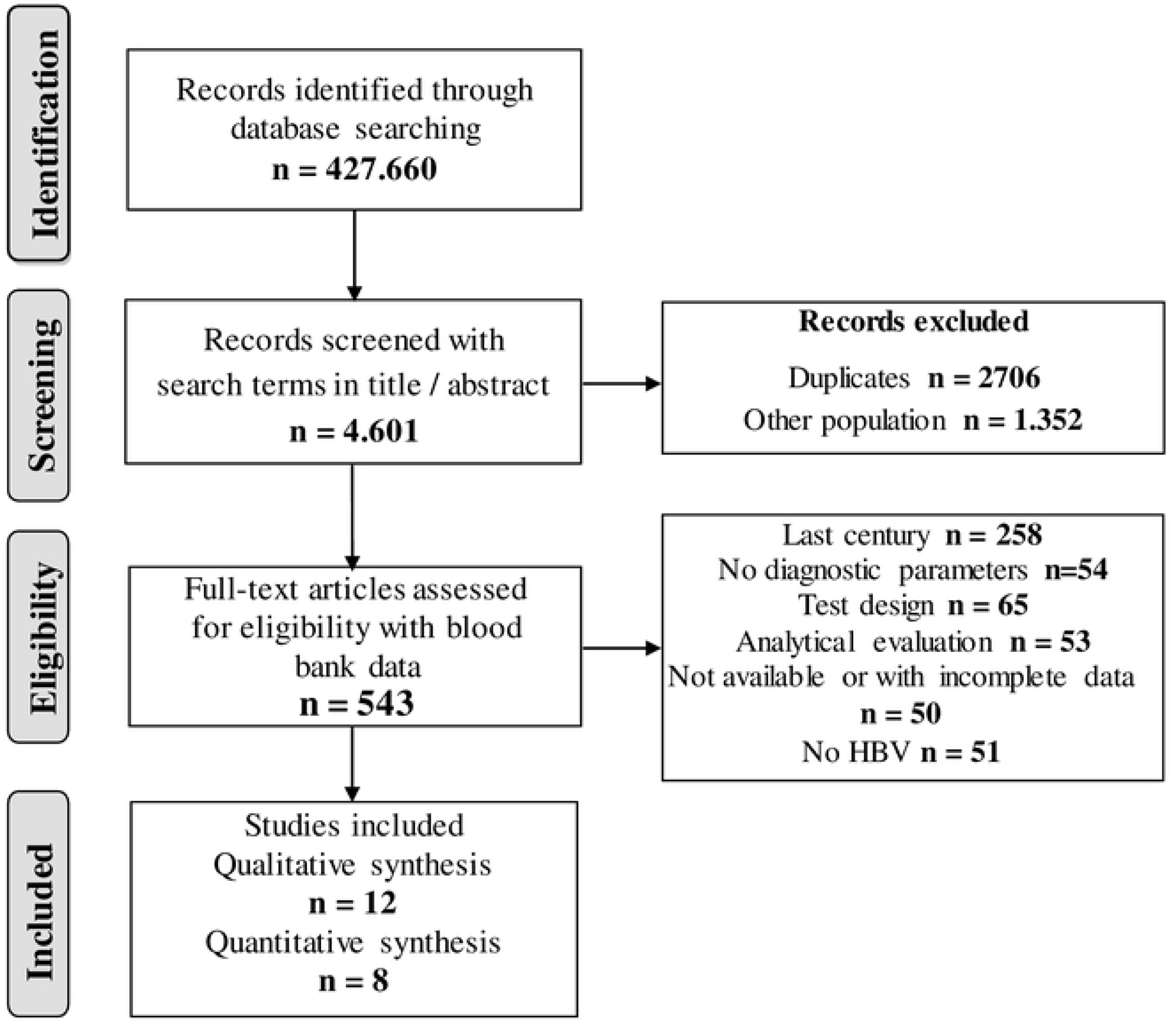
Flowchart of search and research selection.

Only two studies evaluated the diagnostic validity of detecting anti-HBc [3,9] and only one study evaluated both HBsAg and anti-HBc detections simultaneously [3]. The other studies evaluated immunological tests for the detection of HbsAg, using as standard a third-generation ELISA test, and only three studies did the index test with a PCR assay [2,5,9]. All studies presented high methodological quality by meeting more than 90% of the criteria of the QUADAS guide, the least applied were those related to the use of thresholds report of indeterminate results and, the appropriate time between the application of the test and the standard, which were applied in 92% of studies.

Based on eight studies evaluating ELISA for HBsAg detection compared with the combination of other immunodiagnostic tests, 28 subgroups were analyzed in 680 HBV-infected and 1840 healthy donors. It is important to clarify that in Jeremiah et al.’s study, they used ELISA as a standard test to detect anti-HBc-IgM antibodies; however, due to the lack of an adequate comparison group this study was not included in this meta-analysis [18]. Similarly, Farooq et al. and Ha et al. used PCRs as standard and were omitted from meta-analysis due to a lack of a standard comparable to other studies [2,5]. The studies of Schmidt et al. and El-Ghitany et al. could not be analyzed because they only evaluated HBc, with sensitivity results ranging from 85.5% to 99.4%, and specificity results between 98.4% and 99.9% [3,9].

Overall sensitivity of HBsAg was 94.1% (95% CI= 92.9% - 95.1%), ranging from 50.0% (95% CI = 29.1% - 70.9%) and 100.0% (95% CI = 96.5% - 100.0%); while overall specificity was 98.2% (95% CI = 97.8% - 98.6%) with a range between 86.9% (95% CI = 79.0% - 92.7%) and 100.0% (95% CI = 96.6% - 100.0%). No study or subgroup had statistically greater weight on the overall measure, demonstrating the relevance of the value estimate of these parameters in a combined way. Both the I^2^ (>50%) and the Chi-square test (p<0.05) showed the heterogeneity in the studies for the parameters of sensitivity and specificity; this evidenced the need to estimate the combined measure through random effects (considering intra and inter studies variation) (Figure 2).

**Figure 2.** Sensitivity and specificity of the immunological diagnosis with HBsAg. (A) correspond to sensitivity analysis and (B) to specificity analysis with confidence intervals (CI) of 95%.

The authors of studies with the lowest sensitivity indicated that these results are attributed to the deficiency of the tests to detect genotypes and subtypes other than HBV in diverse populations, and also due to technical problems including: lack of quality assurance, poor training, and recertification of laboratory staff in rural areas, problems transporting and storing samples in resource-deficit remote areas, the absence of controls and a lack of evaluation of commercial kits [3,10,17].

The overall positive LR was 53.06(95% CI = 32.38 - 86.96) with a range between 7.4 (95% CI = 4.6 - 12.0) and 2633.8 (95% CI = 164.4 - 42186.3), and the overall negative LR was 0.05 (95% CI = 0.023-0.092), ranging from 0.005 (95% CI = 0.000-0.075) to 0.507 (95% CI = 0.340-0.757), with no study with greater statistic weight on the combined measurement. In these parameters the I^2^ (>50%) and Cochran-Q test (p<0.05) showed the heterogeneity in the studies (Figure 3).

**Figure 3.** HBsAg likelihood ratios. (A) correspond to positive LR and (B) correspond to negative LR both with CI of 95%

The global diagnostic OR was 1720.9 (95% CI = 607.18 - 4418.8), with a range of 34.7 (95% CI = 18.4 - 65.2) to 45365.0 (95% CI = 891.91 - 2307390), and an area under curve of 99.72%; with a sensitivity analysis that demonstrated the robustness of the combined measurement. Also high heterogeneity was found according to I^2^ and Cochran-Q (Figure 4).

**Figure 4.** Odds ratio diagnostic and area under curve for HBsAg detection. (A) Corresponds to ORD and (B) to the Summary Receiver Operating Curve (SROC).

## Discussion

This review included 12 studies with high methodological quality in which different diagnostic tests were applied for the detection of HBV in 18,620 individuals and HBsAg meta-analysis in 2,520 participants. The results confirmed the advantages offered by the meta-analysis of diagnostic tests, allowing extrapolation of their results, greater accuracy, high quality evidence and diagnostic performance. These parameters can be used to improve clinical and epidemiological decision-making [13,20].

The continents with the most research on this topic were Africa and Asia, and in contrast, America and Europe had an inadequate report of diagnostic test evaluations for HBV, which relates to some epidemiological data. For example, the latest World Health Organization (WHO) report for 2019, indicates that the West Pacific (Asian) and African regions had higher HBV infection prevalence (around 6%), while the Eastern Mediterranean, South-East Asia, Europe and the Americas were lower with a 3.3%, 2.0%, 1.6% and 0.7% respectively [21].

The combination of the studies included in this analysis, allowed us to calculate an overall measure for sensitivity and specificity of 94% and 98% respectively, indicating that there are few false negative and positive results in those immunological tests for the detection of HbsAg. In addition, PLR of 53 and NLR of 0.05, confirm that the gain of sensitivity and specificity in the overall measurement of the evaluated tests, is not due to false positive or negative results. The high Odd Radio Diagnostic ORD reflects HBsAg’s excellent ability to differentiate truly healthy from infected individuals, as HBV-infected donors are 1,721 times more likely to generate a positive result, compared to healthy donors (the odds of a false positive are extremely low). Likewise, the ROC curve showed an excellent relationship between sensitivity and specificity with a value of 0.9972; being a statistical parameter that allows different studies to be grouped under the same curve, and confirming that there is a high diagnostic performance in the detection of HBsAg [22]. Only two studies evaluated Anti-HBc, indicating that the use of immunological tests for the detection of HBsAg, remain the most widely used in the screening of blood donors. The HBsAg is the first serological marker that appears after infection with this virus and can even be detected during the incubation period, and its persistence for a period longer than six months allows one to classify the infection as chronic [19]. However, authors such as Schmidt et al. and Jeremiah et al. indicate that detection of anti-HBc should be considered as a potential measure to increase the safety of the blood to be transfused, since HBsAg detection does not rule out the risk of HBV transmission during the immune window period. In addition, the anti-HBc appears first during the acute period, reflecting recent infection, and making it an excellent marker of hidden HBV infection. Also, anti-HBc detection has lower cost compared to PCR, and may be useful in people who conceal risky behaviors in pre-donation surveys [9,18,23].

Finally, these findings account for the validity and safety of HBsAg detection for screening of blood banks, and its potential utility in clinical contexts. This is consistent with the WHO Report on Hepatitis B for 2019, which reiterates its recommendation to test all blood units with this marker in order to detect infection, ensure blood safety, prevent HBV transmission, and improve access to treatment for the infected donors. These findings are of greater importance when considering that in 2016, WHO estimated about 27 million people infected and only 16.7% of the diagnosed population has access to treatment, principally due to the large limitations they have to access diagnosis. The lack of timely diagnosis increases the risk of transmission of the infection and thus the possibility of developing cirrhosis or hepatocellular carcinoma which by 2015 caused 887,000 deaths [21].

Among the main limitations of this meta-analysis is the lack of demographic and clinical-epidemiological data of the participants of each study to analyze possible sources of heterogeneity in the parameters analyzed and perform analyses of Subgroups.

## Conclusion

With a large sample size, for both healthy and infected donors and studies of high methodological quality, it is confirmed that the detection of HBsAg has excellent validity which supports its use in clinical screening, blood banks and population-surveillance programs.

## Supporting information

S1. Search strategies applied to databases.

## Acknowledgments

None apply.

## Author Contributions

Conceptualization: JACA.

Data curation: CPOM, JACA.

Funding acquisition: JCZ.

Investigation: CPOM, JACA, SMM, JCZ.

Methodology: CPOM, JACA.

Software: CPOM, JACA.

Supervision: JACA, JCZ.

Validation: CPOM, JACA, SMM, JCZ.

Writing – original draft: CPOM, JACA, SMM, JCZ

Writing – review & editing: CPOM, JACA, SMM, JCZ.

## Funding

This research received no external funding.

## Conflicts of Interest

None of the authors declare conflict of interest for the publication of this manuscript.

## References

1. World Health Organization. New hepatitis data highlight need for urgent global response. 2017. [Cited 2014 March 17]. Available: https://www.afro.who.int/news/new-hepatitis-data-highlight-need-urgent-global-response.

2. Farooq A, Waheed U, Zaheer HA, Aldakheel F, Alduraywish S, Arshad M. Detection of hbsag mutants in the blood donor population of pakistan. PLoS One. 2017;12(11):e0188066. doi:10.1371/journal.pone.0188066.

3. El-Ghitany EM, Farghaly AG. Evaluation of commercialized rapid diagnostic testing for some hepatitis b biomarkers in an area of intermediate endemicity. J Virol Methods. 2013;194(1-2):190–3. doi: 10.1016/j.jviromet.2013.08.026

4. Randrianirina F, Carod JF, Ratsima E, Chretien JB, Richard V, Talarmin A. Evaluation of the performance of four rapid tests for detection of hepatitis b surface antigen in antananarivo, madagascar. J Virol Methods. 2008;151(2):294–7. doi: 10.1016/j.jviromet.2008.03.019.

5. Ha J, Park Y, Kim HS. Evaluation of clinical sensitivity and specificity of hepatitis b virus (hbv), hepatitis c virus, and human immunodeficiency virus-1 by cobas mpx: Detection of occult hbv infection in an hbv-endemic area. J Clin Virol. 2017;96:60–3. doi: 10.1016/j.jcv.2017.09.010.

6. Dogbe EE, Arthur F. Diagnostic accuracy of blood centers in the screening of blood donors for viral markers. Pan Afr Med J. 2015;20:119. doi: 10.11604/pamj.2015.20.119.5263.

7. Maity S, Nandi S, Biswas S, Sadhukhan SK, Saha MK. Performance and diagnostic usefulness of commercially available enzyme linked immunosorbent assay and rapid kits for detection of hiv, hbv and hcv in india. Virol J. 2012;9:290. doi: 10.1186/1743-422X-9-290.

8. Bedoya JA, Cortes Marquez M, Cardona-Arias JA. Seroprevalence of markers of transfusion transmissible infections in blood bank in colombia. Rev Saude Publica. 2012:46(6):950–9.

9. Schmidt M, Nubling C, Scheiblauer H, Chudy M, Walch LA, Seifried E, et al. Anti-hbc screening of blood donors: A comparison of nine anti-hbc tests. Vox Sang. 2006;91(3): 237–43.

10. Mutocheluh M, Owusu M, Kwofie TB, Akadigo T, Appau, E, Narkwa PW. Risk factors associated with hepatitis b exposure and the reliability of five rapid kits commonly used for screening blood donors in ghana. BMC Res Notes. 2014;7: 873. doi: 10.1186/1756-0500-7-873.

11. Molina M. El metaanálisis de pruebas diagnósticas. Rev Pediatr Aten Primaria, 2015;17(6): 281–5.

12. Zamora J, Abraira V, Muriel A, Khan K, Coomarasamy. A. Meta-disc: A software for meta-analysis of test accuracy data. BMC Med Res Methodol. 2006;6(31). doi:10.1186/1471-2288-6-31.

13. Cardona-Arias J, Ríos L, Higuita-Gutierrez L. Revisiones sistemáticas de la literatura. Bogotá: Univerisdad Cooperativa de Colombia; 2019. DOI: http://dx.doi.org/10.16925/9789587600377.

14. Moher D, Liberati A, Tetzlaff J, Altman DG, Group PRISMA. Preferred reporting items for systematic reviews and meta-analyses: The prisma statement. Open Med. 200;6(7): e1000097. doi: 10.1371/journal.pmed.1000097.

15. Whiting PF, Rutjes AW, Westwood ME, Mallett S, Deeks JJ, Reitsma JB, et al. Quadas-2: A revised tool for the quality assessment of diagnostic accuracy studies. Ann Intern Med. 2011;155(8): 529–36. doi: 10.7326/0003-4819-155-8-201110180-00009.

16. Louisirirotchanakul S, Kanoksinsombat C, O’Charoen R, Fongsatikul L, Puapairoj C, Lulitanond V, et al. Hbsag diagnostic kits in the detection of hepatitis b virus mutation within “a” determinant. Viral Immunol. 2006;19(1): 108–114.

17. Liu S, Figueroa P, Rou K, Wu Z, Chen X, Detels R. Safety of the blood supply in a rural area of china. J Acquir Immune Defic Syndr. 2010; 53(Suppl 1): S23–26. doi: 10.1097/QAI.0b013e3181c7d494.

18. Jeremiah ZA, Idris H, Ajayi BB, Ezimah AC, Malah MB, Baba MM. Isolated anti-hbc-igm antibody among blood donors in the semi-arid region of nigeria. Hum Antibodies. 2011;20(3-4): 77–82. doi: 10.3233/HAB-2011-0242.

19. Kilale AM, Range NS, Ngowi PH, Kahwa AM, Mfinanga SG. Evaluation of the kemri hep-cell ii test kit for detection of hepatitis b surface antigens in tanzania. Tanzan J Health Res. 2012;14(3): 189–193.

20. Beltran O. Revisiones sistemáticas de la literatura. Rev Col Gastroenterol. 2005; 20(1):60-9.

21. World Health Organization. Hepatitis B. 2019. [Cited 2014 March 17]. Available: https://www.who.int/news-room/fact-sheets/detail/hepatitis-b

22. Torres D, Sierra F, Beltran O. Cuando la evidencia evalúa pruebas diagnósticas… ¿qué debemos saber, qué debemos hacer? Rev Col Gastroenterol, 2004;19(4): 281–285.

23. Kumar H, Gupta PK, Jaiprakash M. The role of anti-hbc igm in screening of blood donors. Med J Armed Forces India. 2007;63(4): 350–2. doi: 10.1016/S0377-1237(07)80013-X.

